# Dynamics of spreading microbial swarms and films

**DOI:** 10.1101/344267

**Authors:** Siddarth Srinivasan, C. Nadir Kaplan, L. Mahadevan

## Abstract

Bacterial swarming and biofilm formation are collective multicellular phenomena through which diverse microbial species colonize and spread over water-permeable tissue. During both modes of surface translocation, fluid uptake and transport play a key role in shaping the overall morphology and spreading dynamics. Here, using *Bacillus subtilis* as our model experimental system, we develop a generalized two-phase thin-film model that couples hydrodynamics, mechanics, osmotic flux and nutrient transport to describe the expansion of both highly motile bacterial swarms, and sessile bacterial biofilms. We show that swarm expansion corresponds to steady-state solutions in a nutrient-rich, capillarity dominated regime. In contrast, biofilm colony growth is described by transient solutions associated with a nutrient-limited, extracellular polymer stress driven limit. We apply our unified framework to explain a range of recent experimental observations associated with the shape, form and dynamics of *Escherichia coli* and *Bacillus subtilis* swarms and biofilms. Our results demonstrate how hydrodynamics and transport serve as key physical constraints in regulating biological organization and function in microbial communities.

## I. INTRODUCTION

Bacteria employ sophisticated surface translocation machinery to actively swarm, twitch, glide or slide over solid surfaces [1–4]. Collectively, they also aggregate into multicellular communities on hydrated surfaces and exhibit large-scale coordinated movement [5]. Surface motility in macroscopic colonies occurs primarily via two distinct modes: either by rapid flagella-mediated swarming expansion [6, 7], or alternatively by slow biofilm expansion driven by extracellular polymer matrix production [8]. In both cases, an interplay between mechanical constraints and biological organization sets limits on the overall colony morphology and expansion dynamics [9]. The forces driving colony expansion are generated by non-homogeneous patterns of biological activity, originating from spatial localizations in cell growth and division [10], extracellular polymer matrix production [11–13], osmolyte secretion [14] and active stresses [15, 16]. Conversely, the formation of localized biologically active zones is tightly coupled to the heterogeneity of the environment, including the diffusion and transport of nutrients [17], accumulation of metabolic by-products [18, 19] and presence of quorum sensing and signaling agents that regulate cell-differentiation and development.

Consequently, the dynamics of colony growth requires a mechanistic description that accounts for spatiotemporal inhomogeneities in biological activity, emergent forces, and flows that transport metabolic agents. In bacterial swarming, cells within the colony are actively propelled by the rotation of flagella on a thin layer of fluid extracted from the underlying soft tissue or gel [1]. In contrast, bacterial biofilms are surface aggregates of sessile bacteria embedded in a self-generated extracellular polymer matrix [20]. Despite marked differences in regulatory genetic pathways, morphology and cell function [5], physical characteristics such as the fluidization of the substrate/tissue, gradients in nutrient availability, the low-aspect-ratio geometry and the existence of multiple phases (*i.e.* cells, biopolymer and fluid) are common to both bacterial film and swarm colonies. Motivated by these similarities, we present a unified multiphase framework that couples mechanics, hydrodynamics and transport to explain the dynamics of bacterial swarm and film expansion.

### Significance statement

Collective microbial swarming and biofilm colonization on hydrated surfaces are important in many natural settings. The two expansion modes are typically differentiated under the paradigm of genetic regulation and cellular signaling/competition, often neglecting the critical role of the physical micro-environment. We present a unified multiphase hydrodynamic theory for the dynamics of swarms and biofilms by addressing the fundamental constraints of water and nutrient availability on local biomass growth. Our results provide a quantitative framework for a number of unexplained experimental observations of microbial colony shape, speed, dynamics in both swarms and films, and set the stage for understanding how to control them.

## II. EXPERIMENTAL BACKGROUND

### A. Bacterial swarms

Experiments on swarming colonies of *E. coli* [14, 21, 22], *S. enterica* [23–26] and *P. aeruginosa* [27] reveal certain reproducible features associated with this modality of collective behavior. For example, *E. coli* swarms on agarose gels have a steady front shape that propagates radially at a uniform speed [22]. In these swarms, measurements of the osmotic pressure profiles were found to be consistent with the active secretion of lipopeptides and glycolipids in regions of high cell density that serve to fluidize the swarm by extracting water from the underlying tissue, thus allowing it to spread [14]. These observations are not unique to *E. Coli*; indeed our experiments with *B. subtilis* swarms, following [28], indicate the same phenomena, *i.e.* a steady-state front shape and speed, as shown in Figs. 1A-1E. Close to the spreading front, we observe a multilayer region of width *W* = 195 *µ*m ± 35 *µ*m, indicated by the dashed white lines in Figs. 1B and 1C. The multilayer region correlates with increased colony thickness and local bacterial density [22]. At the edge, and in the interior, there is just a monolayer of cells. The swarm radial expansion velocity is constant at *V* = 2 mm/hr (see Fig. 1D) and the swarm front maintains a steady-state profile during expansion (see Fig. 1E). These observations raise a number of natural questions associated with the steady-state velocity and profile of the swarm colony. Given the observations of osmotic gradient-driven flow in the vicinity of the growing front [14], coupled with variations in the thickness and activity of bacteria, any framework to explain these requires a consideration of a dynamic bacterial population interacting with ambient fluid, necessitating a multiphase description.

**Figure 1.**
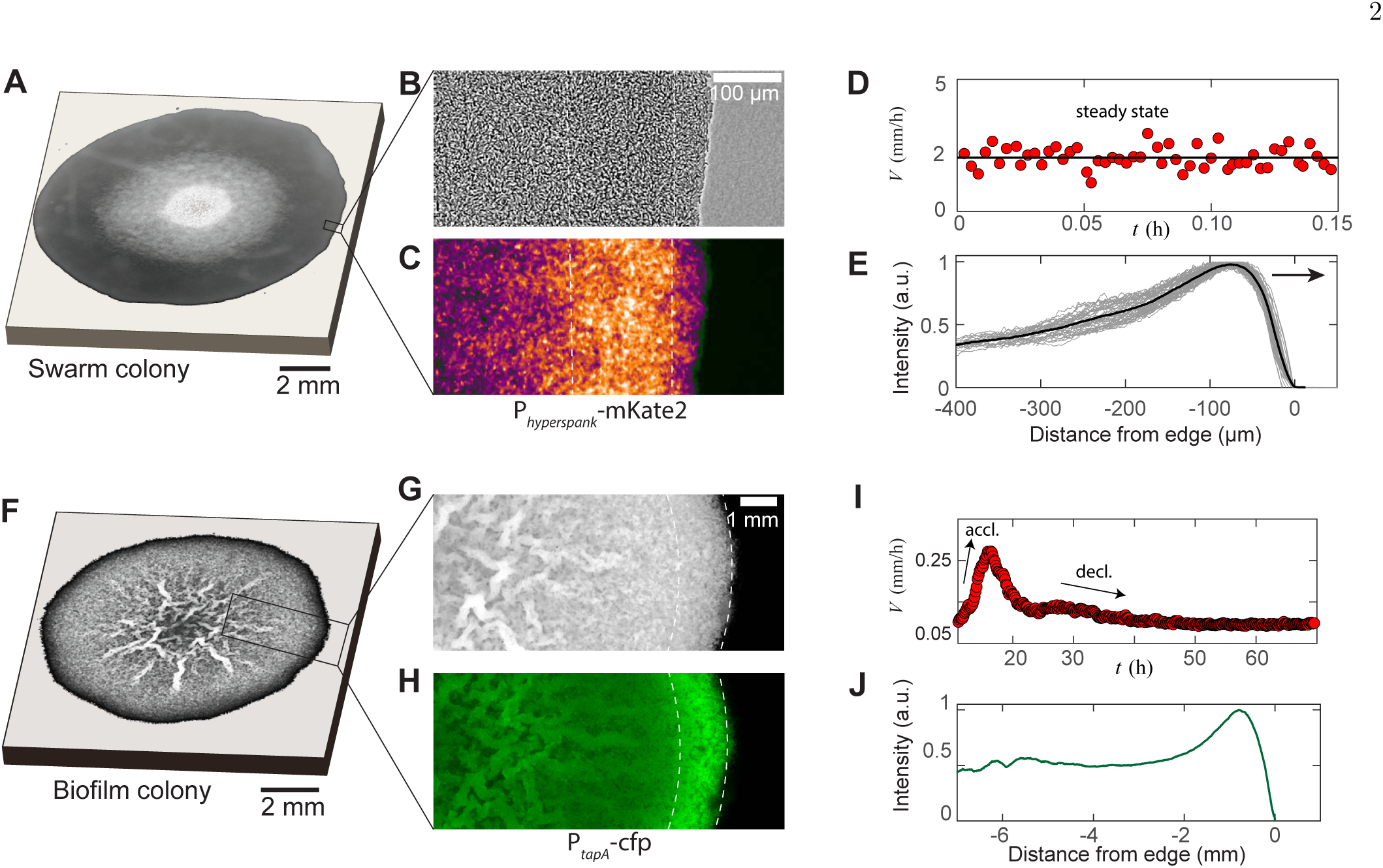
Experimental features of microbial swarms and biofilms. (A) Snapshot of a *Bacillus subtilis* swarm expanding on a 0.5 wt% LB/agar gel. (B,C) Brightfield and fluorescent zoom images of the leading swarm edge of a MTC822 strain containing the fluorescent *P_hyperspank_*-mKate2 reporter that is expressed constitutively. The dashed white lines indicates the extent of the multi-cellular region. (D) Expansion velocity of the swarm measured at intervals of 10 s over a 10 min period. The solid line corresponds to a mean steady-state velocity of *V* = 2 mm/h. (E) Mean intensity traces of the constitutive fluorophore (mKate2) representing bacterial densities profiles plotted in the moving steady-state frame. The dark grey traces represent separate density profile measurements taken every 10 s in the advancing swarm. The solid line represents the density profile averaged over a period of 30 min. (F) A *Bacillus subtilis* biofilm colony developing on a 1.5 wt% MSgg/agar gel. (G,H) Brightfield and fluorescent zoom images of the biofilm colony formed by a MTC832 strain harboring the *P_tapA_*-cfp fluorescent reporter expressed in cells synthesizing the extracellular polymeric matrix (EPS). The dashed white lines indicates the extent of an active peripheral zone signifying localized EPS production. (I) Expansion velocity of the biofilm colony measured at intervals of 10 mins over a 72 h period. The peak expansion velocity of *V* = 0.22 mm/h occurs at *t* ~ 18 h after inoculation. (J) Azimuthally averaged matrix reporter activity (cfp) as a function of spatial distance within the biofilm.

### B. Bacterial biofilms

In contrast with swarms, in spreading bacterial biofilms, the extracellular polymeric substance (EPS) matrix facilitates expansion via osmotic fluid influx in *B. subtilis* [11] and *V. cholerae* [12] biofilm colonies. However, EPS synthesis is not homogeneous, and depends on the local nutrient concentration and environmental heterogeneities experienced by cells within the same biofilm [29, 30]. Recently, it was shown that the EPS matrix production is localized to cells in the propagating front of the biofilm [13]. Repeating these experiments, we focus on a peripheral region of a biofilm colony using a *B. subtilis* strain (MTC832) that harbors the *P_tapA_* − cfp construct as a reporter for matrix production activity [13, 31], as shown in Figs. 1F-1J. This highlights a ~ 1 mm zone of matrix production activity at the periphery, seen in Fig. 1G and 1H; indeed plots of averaged matrix production reporter intensity exhibit a distinct peak at the periphery, as shown in Fig. 1J. Looking at the dynamics of radial expansion following Ref.[13] confirms the existence of an initial acceleration regime followed by an abrupt transition to a second regime characterized by a monotonic decrease in expansion velocity, as plotted in Fig. 1I. This transient mode of biofilm spreading driven by EPS production and swelling is quite different from that of bacterial swarming, and suggests that we need a fundamentally different way to address its origins. However, if we now consider the EPS matrix and fluid as distinct phases [11, 32–34], with the bacterial population being relatively small, we are again led to a multiphase description of the system, but with a different dominant balance relative to that seen in bacterial swarms.

## III. THEORETICAL FRAMEWORK

Recent theoretical approaches have considered specific physical factors such as the wettability of the biofilm [35, 36], osmotic pressure in the EPS matrix [11, 34], or Marangoni stresses associated with the swarm fluid [37], in an attempt to rationalize the dynamics of colony expansion. However, a description that captures the experimental observations described in Fig. 1 remains lacking. Here, given the similarities between the bacterial swarming and filming systems, we provide a unified description of their spreading dynamics by recognizing that in both cases we need to consider large slender microbial colonies with *h*^∗^*/R* ≪ 1, where *h*^∗^ is the colony thickness and *R* is the radius. This approximation results in a quasi-2-dimensional, two-phase model (assuming axisymmetry) of a colony that spreads along the *x*-axis, with a varying thickness, as shown in Fig. 2. The subscript *i* = (1, 2) denotes the actively growing phase and passive phase, respectively. Within the swarm colonies, the highly motile cells constitute the actively growing phase whereas the fluid comprises the passive phase. Similarly, in biofilms, the EPS matrix constitutes the active phase, and the aqueous fluid is the passive phase. In both cases, colony growth occurs over a semipermeable soft gel/tissue substrate, as shown in Fig. 2. We develop a continuous description of colony expansion in terms of averaged variables associated with the position of the upper colony interface, *h*(*x, t*), the volume fraction of the active phase (*i.e.*, swarmer cells or polymer matrix) *ϕ*_1_ = *ϕ*(*x, t*) while the volume fraction of the fluid phase is *ϕ*_2_ = 1 − *ϕ*(*x, t*), which are coarse-grained variables extracted from an averaging procedure over local fluctuations [38, 39]. The 1-D substrate depth-averaged nutrient concentration field within the substrate is *c*(*x, t*). As detailed in Appendix [B], combining mass and momentum balances yields the following generalized set of partial differential equations that governs the dynamics of both expanding swarms and biofilms,

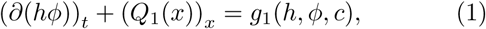

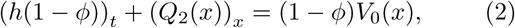

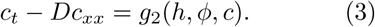

where, (·)*_x_* = *∂* (·) */∂x*, etc. Here, *Q*_1_(*x*) is the horizontal flux in the active phase, *Q*_2_(*x*) is the horizontal flux in the fluid phase, and *V*_0_(*x*) is the osmotically-driven net vertical fluid influx per unit length across the permeable substrate. The dynamics of swarms and biofilms differ in the details of the expressions for *Q*_1_*, Q*_2_, *V*_0_ are provided in Table. I, and discussed in the following sections.

**Figure 2.**
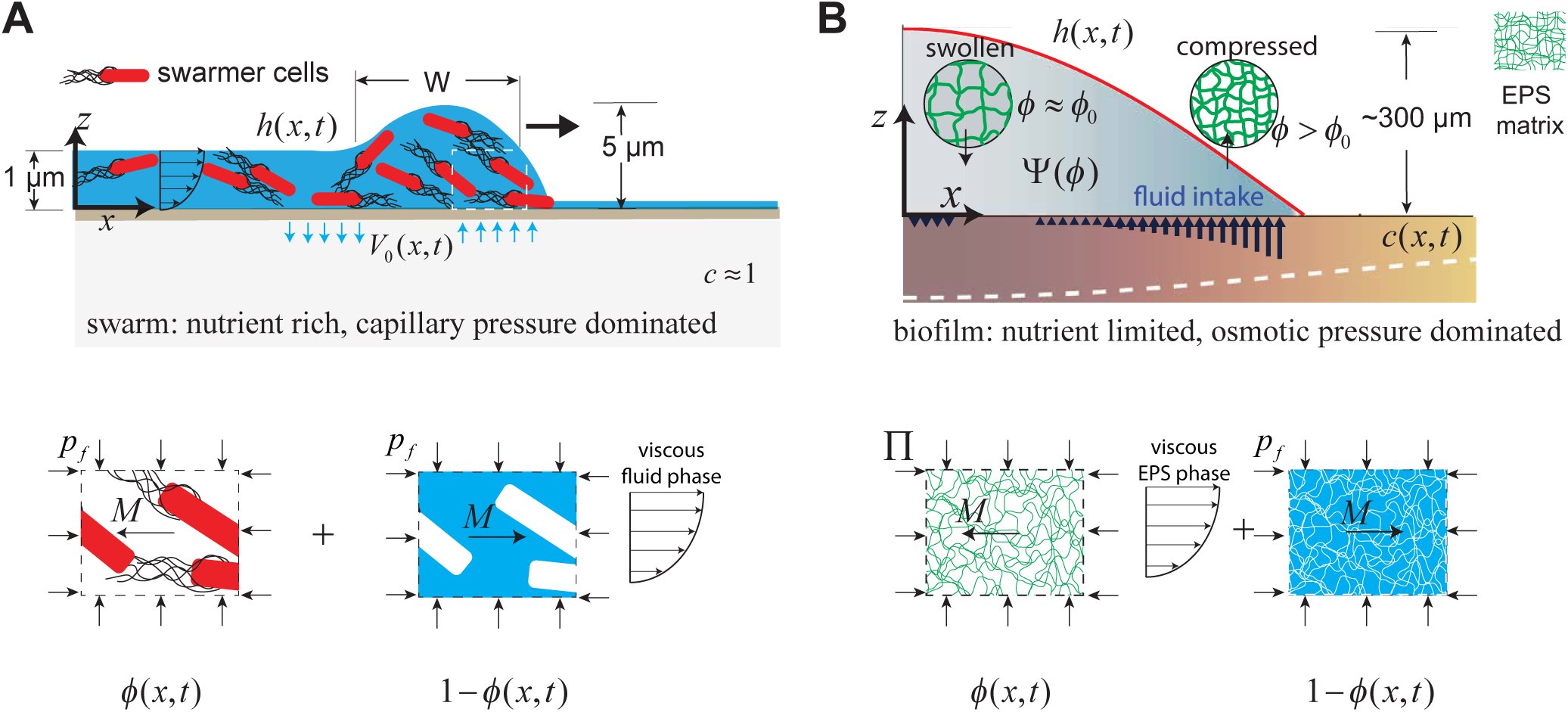
Geometry and variables governing colony expansion in (A) microbial swarms, and (B) bacterial biofilms, respectively. In both cases, the total thickness of the microbial colony is *h*(*x, t*), the averaged nutrient concentration field is *c*(*x, t*), the volume fraction of the active phase is *ø*(*x, t*), the volume fraction of the fluid phase is 1 − *ø*(*x, t*), and the fluid influx across the agar/colony interface is denoted by *V*_0_(*x, t*). As shown on the bottom panel, the active phase constitutes swarmer cells in the microbial swarm, and secreted EPS polymer matrix in the biofilm. The pressure in the fluid phase is *p_f_* and the effective averaged pressure in the active phase is Π. In the swarm cell phase, Π = *p_f_*, while the EPS phase effective pressure is Π = *p_f_* + *ø*Ψ(*ø*), where Ψ(*ø*) is swelling pressure and is related to Flory-Huggins osmotic polymer stress. The momentum exchange between the two phases is denoted by *M*, which includes the sum of an interfacial drag term and an interphase term as detailed in Eq. (B11) in the Appendix.

**Table I.** Definitions of the active phase horizontal flux *Q*_1_, the fluid phase horizontal flux *Q*_2_, active phase growth term *g*_1_(*h, ø, c*), osmotic influx term *V*_0_(*x*), and nutrient consumption term *g*_2_(*h, ø, c*) for bacterial swarms and films in the generalized thin film evolution equations described by Eqs. (1)-(3). Here, *µ*_1_ is the biofilm viscosity, *µ*_2_ is the fluid viscosity, *pf* is the fluid phase pressure, Π is the effective pressure in the active phase, *g*_0_ is effective swarmer cell growth rate, *G* is the EPS production rate, Γ is the nutrient consumption rate per unit concentration, *K* is the nutrient half-velocity constant and *d* is the thickness of the substrate. For swarms, the active phase corresponds to the swarmer cell phase, and for biofilms, the active phase is the EPS polymer matrix.

|  | Variables | Swarms | Biofilms |
| --- | --- | --- | --- |
| Flux<br>(Phase I) | $Q_1(x)$ | $-\frac{h^3}{3\mu_2} \frac{\phi}{1-\phi} \frac{\partial p_f}{\partial x} - \frac{h\phi}{\zeta} \frac{\partial p_f}{\partial x}$ | $-\frac{h^3}{3\mu_1} \frac{\partial \Pi}{\partial x}$ |
| Flux<br>(Phase II) | $Q_2(x)$ | $-\frac{h^3}{3\mu_2} \frac{\partial p_f}{\partial x}$ | $-\frac{h^3}{3\mu_1} \frac{1-\phi}{\phi} \frac{\partial \Pi}{\partial x} - \frac{h}{\zeta} \frac{(1-\phi)^2}{\phi} \frac{\partial p_f}{\partial x}$ |
| Osmotic influx | $V_0(x)$ | $Q_0 \left( \frac{\phi}{1-\phi} - \frac{\phi_0}{1-\phi_0} \right)$ | $Q_0 (\phi^3 - \phi_0^3)$ |
| Growth term | $g_1(h, c, \phi)$ | $g_0 h \phi \left( 1 - \frac{h\phi}{h^* \phi_0} \right)$ | $G h \phi \frac{c}{K+c} \left( 1 - \frac{h\phi}{h^* \phi_0} \right)$ |
| Nutrient uptake | $g_2(h, c, \phi)$ | - | $\frac{\Gamma \phi h}{d} \frac{c}{K+c}$ |

## IV. MICROBIAL SWARMS

Existing thin-film models to describe bacterial swarming assume that gradients in wetting agent activity generate Marangoni stresses that drives swarming motility [37, 40]. However, recent experiments [27] demonstrates that *P. aeruginosa* swarms robustly even after exogenously eliminating gradients in surfactant concentration within the swarm fluid, eliminating Marangoni flows as the principal mechanism that drives swarming. Here, we take a different approach based on experiments that show that steady-state swarm colony expansion is mediated by osmolyte secretion [22]. As we will see, this leads to fluid being extracted from the substrate near the front, then driven into the colony by capillary and viscous stresses, and eventually returns into the substrate in the interior of the swarm.

Within the bacterial swarms, the dominant phases are the swarmer cell phase, and the viscous aqueous phase, as shown in the bottom panel of Fig. 2A. Fluid uptake from the substrate is regulated by the secretion of osmotically active agents by the swarmer cells [14]. We represent the osmotic agent in the fluid by a concentration field, *c*_osm_(*φ*) that is proportional to the local volume fraction of cells such that *c* ∝ *ø/*(1 − *ø*), and gives rise to an osmotic pressure described by the Van’t Hoff’s law as [41], ΔΨ = (Ψ_0_*ø/*(1 − *ø*) − Ψ_eq_), that drives the fluid intake. Here, Ψ_0_ is the osmotic pressure scale in the swarm fluid and Ψ_eq_ is the equilibrium osmotic pressure within the underlying tissue/gel substrate. Away from the front, in the interior of the swarm colony, there is no net fluid in-flux [14]. Therefore, the equilibirium volume fraction of the swarm cells at the interior is, *ø*_0_ = Ψ_eq_*/*(Ψ_0_ + Ψ_eq_). At the front itself, the difference in osmotic pressure results in a net Darcy-type fluid influx into the swarm, *V*_0_(*x*), expressed as,

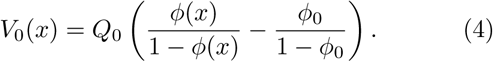

where *Q*_0_ is a velocity scale associated with fluid inflow from the substrate. Measurements of cell replication within swarms reveals that growth is restricted to swarmer cells at the periphery [10], which we model using a modified logistic growth term *g*_1_(*h, φ*) as listed in Table. I, that localizes all cell division to the periphery. Here, *h*^∗^*ø*_0_ is the limiting thickness of the swarmer-cell phase at the interior, and *g*_0_ is an effective specific growth rate, related to true specific cell growth rate by a geometric factor (see discussion in Appendix. [B 1]).

### A. Dimensionless scales and parameters governing bacterial swarms

To make sense of the scales in the problem, we use the dimensionless variables 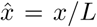, 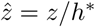 and 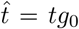 where *h*^∗^ is the vertical length scale, *L* is a horizontal length scale and 1*/g*_0_ is the time-scale associated with bacterial growth. The resultant horizontal velocity scale in the swarm colony is *U* = *Lg*_0_. Swarm expansion is fluid driven, and therefore balancing the viscous stresses generated in the swarm fluid, with the curvature pressure due to surface tension results in *µ*_2_*U/h*^∗2^ ~ *γh*^∗^*/L*^3^, where *µ*_2_ is the viscosity and *γ* is the surface tension of the aqueous phase. As a result, the natural horizontal length scale is *L* ~ *h*^∗^(*Ca*)^−1*/*3^, where *Ca* = (*µ*_2_*U/γ*) is a capillary number associated with the microbial swarm fluid. We define a dimensionless parameter *α*_1_ = (*γh*^∗^*/L*^2^)*/*(*ζLU*) that relates the magnitude of capillary forces to the viscous drag acting on cells within the swarm. Here, *ζ* = *ζ_c_/V_c_* where *ζ_c_* is the friction coefficient of a single swarmer cell and *V_c_* is its volume. Finally, we define *α*_2_ = *Q*_0_*/*(*h*^∗^*g*_0_) as a dimensionless parameter that sets the ratio of fluid influx velocity to a thickness velocity scale associated with bacterial growth.

The vertical length scale and equilibrium fluid volume fraction are estimated from the interior monolayer region as *h*^∗^ = 0.5 *µ*m and *φ*_eq_ = 0.5 [22]. The time scale is determined by choice of the effective growth rate *g*_0_ = 0.013 s^−1^. Experimental measurements of spatial profiles of cell division in swarm colonies remain lacking [10]. Consequently, we have chosen *g*_0_ as a fitting parameter in our study, as discussed in Appendix [B 1]. We assume values of *µ*_2_ = 10^−3^ Pa.s, and *γ* = 10^−2^ N/m leading to a horizontal length scale of *L* = 100 *µ*m, velocity scale of *U* = 1.3 *µ*m s^−1^ and *Ca* = 10^−7^. For cells of diameter *a* = 1 *µ*m, we estimate *ζ_c_* ≈ 0.01 pN/(*µ*m s^−1^), resulting in *α*_1_ ≈ 0.2. The vertical fluid influx velocity scale is set as *Q*_0_ = 10^−2^ *µ*m/s, which corresponds to a value of *α*_2_ ≈ 1.5.

### B. Steady state swarms

Under these scaling assumptions, and considering that nutrient concentration is constant, Eqs. (1)-(3) reduce to the following scaled equations in the swarming limit,

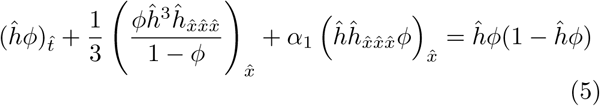

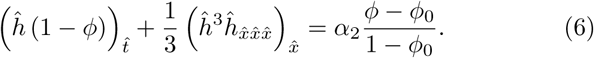

Eq. (5) assumes that bacterial growth is solely fueled by the consumption of an abundant nutrient field. To complete the formulation of the problem, we need five boundary conditions which are 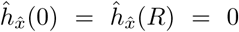, 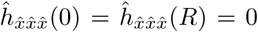, and *ø*(0) = *ø*0, where 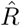 is set much larger than the colony size 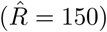 in our simulations. The initial condition corresponds to a circularly inoculated swarm colony, along with a thin pre-wetting film where no bacterial growth occurs (see Fig. C.1).

Solving Eqs. (5)-(6) with the prescribed initial and boundary conditions numerically results in a steady state solution that advances at a constant speed (see Fig. C.2). In Fig. 3, we plot a representative steady state solution in the frame of the advancing front for *α*_1_ = 0.2, *α*_2_ = 1.5 and *φ*_0_ = 0.5. Near the leading edge of the swarm, there is a region of enhanced thickness as indicated by the red line in Fig. 3A, while right behind it, in the multi-cellular region the average cell volume fraction is *ø* ≈ *ø*_0_. Just behind the leading edge, where the cell concentration is highest, so is the osmolyte concentration leading to fluid extraction from the substrate, while further behind, fluid is pulled back in, as indicated by the arrows in Fig. 3A. In Fig. 3B, we show the steady-state osmotic flow solution and see that it correlates well with the experimentally measured osmotic pressure profile by Ping *et al* [14] in *E. coli* swarms. As suggested by earlier experiments [14], this zone of cellular and osmolyte activity near the leading edge drives the advancing swarm front, and is analogous to a capillary ridge in thin film fluid flows with a width *W* ~ *Ca*^−1*/*3^. As shown in Fig. C.3, our numerical horizontal flow profiles are also consistent the scaled radial fluid velocity measurements of Wu & Berg [22]. In Fig. 3C, we see that the radial expansion velocity scales as *h*^∗^*g*_0_ and shows quantitative agreement with experiments and is insensitive to the fluid influx velocity scale when *Q*_0_ ≫ *g*_0_*h*^∗^. This leads to a picture wherein the combination of the fluid-filled substrate and swarm front work together like a localized active circulatory system.

**Figure 3.**
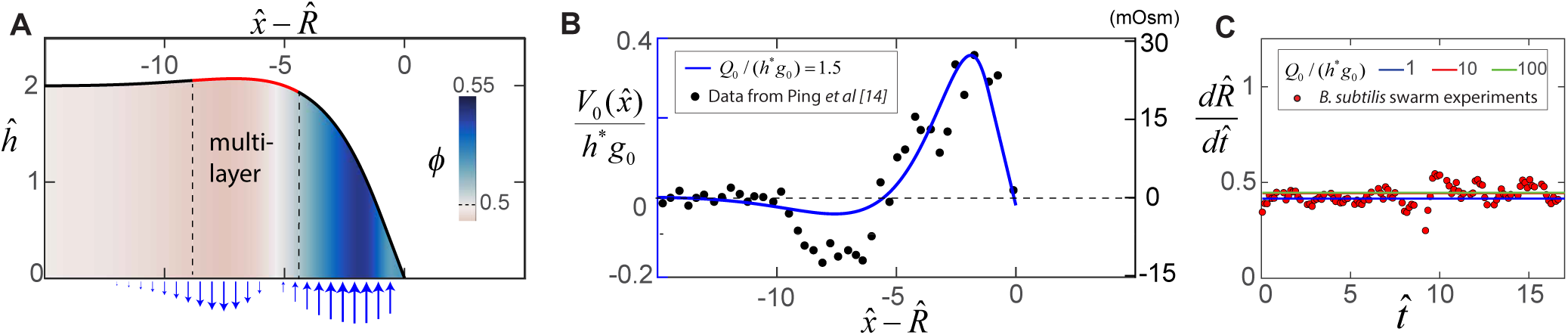
Steady-state morphology and fluid transport in a bacterial swarm obtained by solving (5) and (6) for *α*_1_ = 0.2, *α*_2_ = 1.5 and *ø*_eq_ = 0.5. (A) Plot of the steady-state thickness 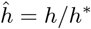 against the scaled distance 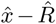, where *x/*(*h*^∗^*Ca*^−1*/*3^) and 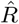 is the radius. The solid red line indicates the extent of the multi-cellular region, and the colormap quantifies variations in *ø*, the local volume fraction. (B) Plotted on the left-axis is the numerical steady-state fluid uptake profile within the swarm (solid line) calculated from (4). On the right axis are experimental measurements of the steady-state osmotic pressure within an expanding *E. coli* swarm (filled circles), reproduced from Ref. [14], with the baseline reference value shifted to zero, and with distances normalized by *L* = 50 *µ*m. (C) Predicted steady-state radial colony expansion speeds within the swarm for values of *α*_2_ = *Q*_0_*/*(*h*^∗^*g*_0_) = 1, 10 and 100 respectively. The data points are expansion speeds in *B. subtilis* swarms measured over 20 minutes, and scaled using *U* = 1.3 *µ*m*s*^−1^ and *g*_0_ = 0.013 s^−1^.

## V. MICROBIAL FILMS

In bacterial biofilms, the EPS matrix secreted by bacteria constitutes the active phase as they swell a lot, drawing in the fluid that acts as the passive phase. As shown in Fig. 2B, the EPS is initially synthesized in a partially swollen, out-of-equilibrium state at the periphery. The polymer chains gradually relax to an equilibrium fully-swollen configuration by the generation of a swelling pressure Ψ within the biofilm, and via fluid uptake *V*_0_(*x*) from the substrate. As discussed in Appendix [B 2], the swelling pressure is Ψ(*ø*) = *ψ*(*ø*)*/ø*, where *ψ*(*φ*) = *ψ*_0_ × *φ*^3^ is the osmotic pressure in the EPS matrix using the Flory-Huggins model for a polymer network in a *θ*-solvent [42], where *ψ*_0_ = *kT /*(*b*^3^) is the osmotic pressure scale, *b* is the approximate size of the monomer unit. The net effective pressure term driving biofilm expansion is, Π = *ψ*_0_*ø*^3^ + *p_f_*, where *p_f_* is the hydrostatic pressure, so that the water influx across the substrate is

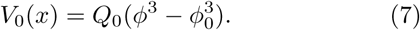

where *Q*_0_ is the influx fluid velocity scale, *φ*_0_ = (Ψ_eq_*/*Ψ_0_)^1*/*3^ is the fully-swollen EPS polymer volume fraction and Ψ_eq_ is the osmotic pressure of the substrate over which the colony grows. Finally, nutrient uptake is modeled by a Monod growth law, while the synthesis of the EPS matrix is modeled by a logistic term as listed in Table I.

### A. Dimensionless scales and parameters in bacterial biofilms

We consider dimensionless variables 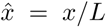, 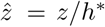, 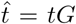, 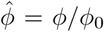 and 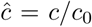, where *h*^∗^ is now the maximum biofilm thickness, *G* is the rate of EPS production, and *c*_0_ is the initial nutrient concentration in the substrate. As biofilm growth is nutrient limited [18], the dimensionless length scale is determined from Eq. (3) as *L* = (*D/G*)^1*/*2^ and the corresponding velocity scale is *U* = (*DG*)^1*/*2^. Using these scales, we can define the ratio of osmotic stresses relative to viscous stress in the EPS phase in terms of the dimensionless parameter, *β*_1_ = (Ψ_0_*/L*)/ (*µ*_1_*U/h*^∗2^), the ratio of capillary other parameter, *β*_2_ = (*γh*^∗^*/L*^3^)/(*µ*1*U/h*∗2), the ratio of capillary stress to the interfacial drag in the aqueous fluid phase, *β*_3_ = (*γh*∗*/L*2)/(*ζU L*), and the ratio of the fluid influx velocity to the EPS swelling velocity, *β*_4_ = *Q*_0_/(*h*^∗^*G*). As shown in Appendix [B], the effective nutrient uptake rate is *S* = (Γ*h*^∗^*ø*_0_)*/*(*c*_0_*d*), where Γ is the nutrient consumption rate per unit concentration and *d* is the substrate thickness. Consequently, we define *β*_5_ = *S/G* as the ratio of the effective nutrient uptake rate to the EPS production rate.

We set the EPS production time-scale as *G* = 1*/*40 min^−1^, resulting in a horizontal length scale of *L* = 1.1 mm and velocity scale *U* = 0.5 *µ*m/s. The effective nutrient uptake rate is estimated as *S* = 1*/*25 min^−1^, where we have taken *d* = 7 mm as the substrate thickness [13], Γ = 10^−2^ mM/s as the nutrient uptake rate [43], and *c*_0_ = 35 mM as the initial concentration of the carbon source. The friction coefficient is *ζ* ~ *µ*_2_*/ξ*^2^, where the EPS mesh size is *ξ* = 50 nm [12]. Using measured estimates of the biofilm viscosity *µ*_1_ = 10^5^ Pa.s [44, 45], fluid phase viscosity *µ*_2_ = 10^−3^ Pa.s, surface tension *γ* = 10^−2^ N/m, an osmotic scale Ψ_0_ = 2100 Pa [12] (*i.e.*, *φ*_0_ = 0.04), biofilm thickness *h*^∗^ = 400 *µ*m, and nutrient diffusivity in agarose gels of *D* = 5 × 10^−10^ *m*^2^/s [43] implies that *β*_1_ ≈ 7, *β*_2_ ≈ 0.01, *β*_3_ ≈ 0.02, *β*_4_ ≈ 1 and *β*_5_ ≈ 2. Consequently, within the context of our model, it is evident that osmotic stresses, fluid influx and biomass growth are the dominant forces that drive colony expansion.

### B. Transient biofilm solutions

With the above scaling assumptions, Eqs. (1)-(3) now reduces to the following partial differential equations that describe biofilm colony expansion,

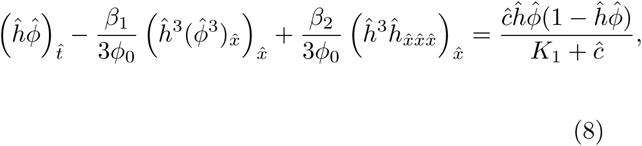

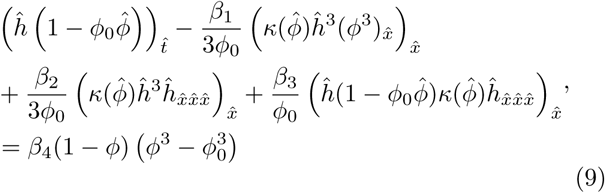

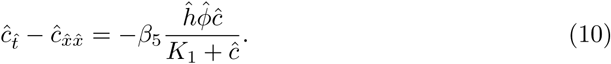

where 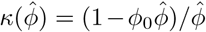 is a volume fraction dependent permeability term. The eight boundary conditions associated with Eqs. (8)-(10) are the symmetry boundary conditions 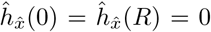, 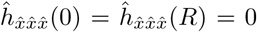, 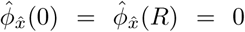, 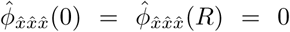 and 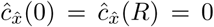, where 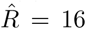 is chosen to match the size of typical 35 mm diameter petri dishes used in experiments [13]. In Fig. 4A, we plot the time evolution of the shape and nutrient concentration field for a biofilm colony of initial radius 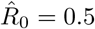 and thickness 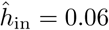.

**Figure 4.**
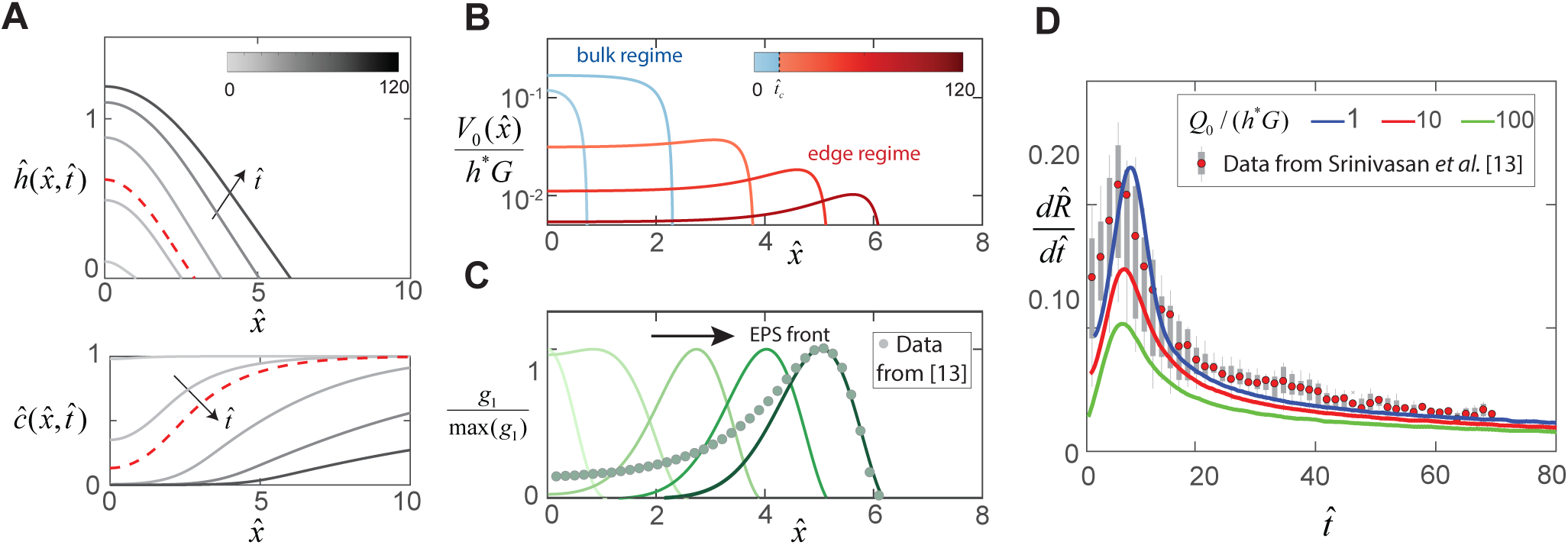
Dynamics of EPS production and biofilm expansion obtained by solving (8)-(10) with *β*_1_ = 6.7, *β*_2_ = 0.01, *β*_3_ = 0.02, *β*_4_ = 1 and *β*_5_ = 1.7. (A) On the top are thickness profiles 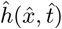 of an expanding biofilm colony, at time intervals of 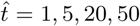 and 90. The nutrient field *c*(*x, t*) at corresponding time intervals is plotted at the bottom. The dotted red line indicates profiles at 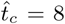, the transition point between the bulk and edge expansion regimes. (B) Variation of the vertical fluid uptake profile within the swarm calculated from (7). The light blue lines correspond to the bulk growth regime for 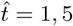 while the red lines correspond to 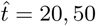 and 90 in the edge growth regime. (C) Plots of normalized EPS production activity within the biofilm, where *g*_1_ is evaluated using the expression in Table. I. The data points are spatial measurements of *tapA* gene activity in *B. subtilis* biofilms reproduced from Ref. [13], with distances scaled by *L* = 550*µ*m. (D) Solid lines indicate transient colony edge expansion velocities for *β*_4_ = 1, 10 and 100 respectively, and with other parameter values fixed as listed above. The experimental data is reproduced from [13] and indicates median expansion velocities (filled circles), the 25th to 75th percentile velocities (filled box), and extreme values (vertical lines), where the data has been scaled by *U* = 0.5 *µ*m*s*^−1^ and *G* = 1*/*40 min^−1^.

Unlike in the case of swarms, the solutions to Eqs. (8)-(10) are transient, and exhibit two distinct expansion regimes: initial acceleration phase until 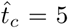, followed by a decelerating phase beyond. For 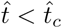, colony expansion arises as the microbes rapidly consumes locally available nutrient at the interior and synthesize fresh EPS matrix, generating spatial gradients in nutrient availability (see Fig. 4A). In Fig. 4B, we show that the newly synthesized EPS generates a large osmotic pressure differential between the biofilm and the substrate, and osmotic fluid influx gradually relaxes the biofilm matrix to a swollen configuration. For *t > t_c_*, the localized zone of EPS production near the film front propagates with a fixed shape as shown in Fig. 4C, consistent with the observed spatial localization in *tapA* gene activity (see Fig. 1J and [13]). Moreover, the radial colony expansion profile in Fig. 4D is also consistent with the non-monotonic front speed observed experimentally [13]. These results are hallmarks of a transition from a bulk to an edge biofilm growth mode, triggered by nutrient limitation. In the latter regime, diffusive transport of nutrients from a region external to the colony continues to sustain EPS production at the biofilm periphery, analogous to Stefan-like, problems in solidification. Our model is thus able to quantitatively rationalize the expansion curves, transition time and localized biological activity observed experimentally, and demonstrates that nutrient availability and diffusive transport governs the dynamics of biofilm growth.

## VI. CONCLUSIONS

Analysis of collective microbial expansion in thin film geometries often prioritizes biological mechanisms, such as genetic regulation, developmental programs and cellular signaling/competition, over the role of the heterogeneous mechanical and physical micro-environments. Here, we have demonstrated that the spatiotemporal dynamics of microbial colony expansion is in fact, primarily governed by the fundamental physical constraints of water and nutrient availability. Using a single generalized multiphase hydrodynamic framework, our model reveals that bacterial swarm colonization corresponds to a fluid regulated limit, whereas the dynamics of biofilm expansion is governed by nutrient transport. In swarms, exudation of water from the permeable substrate via bacterial osmolyte secretion facilitates steady state colony expansion. Numerical solutions of our model demonstrate that the shape of the swarm front is determined by capillarity, and its expansion speed by cell-division and growth. In contrast, transient biofilm expansion is driven by osmotic polymer stresses generated via EPS matrix production in a spatially localized zone at the periphery. Nutrient transport and depletion leads to the formation of these heterogenous zones, and results in two regimes in biofilm expansion. Our results naturally raise the question of controlling biofilm expansion by manipulating water and nutrient availability just as much as manipulating the genetic regulation of EPS production, cell division, and mechanisms of competition and cooperation in microbial colonies.

## Appendix A: Experimental

### 1. Strains

In this study, we used two *B. subtilis* strains, MTC822 and MTC832, that were both previously constructed from a wild-type NCIB 3610 *B. subtilis* strain using a standard transformation protocol [46]. The MTC822 strain is used for fluorescence visualization in the swarming experiment, where the mkate2 red fluorescent protein reports on the activity of the constitutive hyperspank promotor via the *amyE::Phyperspank-mkate2* construct. The MTC832 strain was used in the biofilm experiments in order to visualize localized matrix production activity and harbors the *amyE::PtapA-cfp* construct. In the MTC832 strain, the cfp cyan fluorescent protein reports on the activity of the *tapA* gene that is associated with exopolysaccharide production activity.

### 2. Materials and Methods

Swarm plates were prepared using 0.5 wt% agarose gel (A1296, Sigma) infused with 25 ml of Luria-Bertani (Miller) medium (*i.e.* 10g/L Tryptone 10g/L NaCl 5g/L Yeast Extract, Sigma) and 25 *µ*g/ml Chloramphenicol. Biofilm plates were prepared using 1.5 wt% agarose gel (A1296, Sigma) infused with the standard MSgg biofilm-inducing growth medium [47] (*i.e.* 50 *µ*M MnCl_2_, 5 mM KH_2_PO_4_, 1 *µ*M ZnCl_2_, 50 *µ*M FeCl_3_, 2 mM MgCl_2_, 700 *µ*M CaCl_2_, 50 *µ*g/ml threonine, 50 *µ*g/ml tryptophan, 50 *µ*g/ml phenylalanine, 0.5 wt% glutamate, 0.5 wt% glycerol, 2 *µ*M thiamine and 100 mM MOPS (pH 7)) and 50 *µ*g/ml Spectinomycin. Note that all plates underwent an identical drying protocol prior to use. Freshly poured plates were initially dried with the lid open under a laminar flow hood for 15 minutes. Subsequently, the lid was closed and the dish was cooled at 25 C overnight for a period of 10 hours. All strains were initially grown in fresh Luria-Bertani (Miller) broth medium (Sigma) until mid-exponential phase in a shaker/incubator at 37C. The cultures were diluted to *OD*_650_ = 0.1 and ~ 1*µ*l drop was deposited onto the corresponding swarm (for MTC822) or biofilm (for MTC832) plates. The petri-plates were transfered to a 30C incubator chamber during growth. Fluorescence imaging was performed using a Zeiss Axiozoom.V16 microscope with a PlanNeoFluar Z 1.0x objective (NA 0.25), with a Zeiss 63 HE filter to image the red mkate2 protein, and a Zeiss 47 HE filter to image the cyan cfp protein. For swarm profile measurements, images of the advancing swarm front were captured every 10 seconds over a period of 10 minutes. For biofilm colonies, expansion velocities were measured every 10 minutes over a period of 72 hours following the protocol described in Ref. [13].

## Appendix B: Governing equations

Consider a quasi-2D expanding swarm or biofilm colony where, *x* denotes the horizontal direction and *z* denotes the vertical (thickness) direction. The macroscopic colony is described by (i) the thickness field *h*(*x, t*), (ii) the volume fraction of the active phase (*i.e.* swarmer cells or EPS matrix *ø*(*x, t*) and (iii) the averaged concentration field in the substrate *c*(*x, t*).

### a. Nutrient transport

We assume that 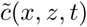 is the nutrient concentration in the substrate denoted by region −*d < z <* 0, where *z* = 0 is the substrate/colony interface and *d* is the substrate thickness. The time evolution of 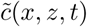 is governed by the diffusion equation.

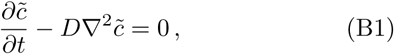

where *D* is the diffusivity of the nutrient in the agarose gel. Integrating the diffusion equation through the substrate thickness results in,

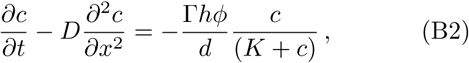

where 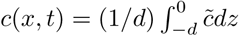 is the mean depth-averaged nutrient concentration in the substrate, Γ is the specific nutrient consumption rate per unit concentration, and *K* is the half velocity constant at which the specific growth rate is one half the maximum value. In obtaining Eq. (B2), we have balanced nutrient efflux from the substrate with nutrient consumption within the colony as 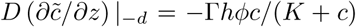. Eq. (B2) describes nutrient transport within the permeable tissue/gel substrate underneath the colony, and holds when the diffusion time scale is much smaller than the flow time scale within the low-aspect-ratio substrate. Upon nondimensionalizing Eq. (B2), the effective nutrient consumption rate *S* used in Table. 1 in the main text is given by *S* = Γ*h*^∗^*ø*_eq_*/*(*c*_0_*d*), where *h*∗ is the colony thickness scale, *φ*_eq_ is the active phase volume fraction scale and *c*_0_ is the initial nutrient concentration in the substrate.

### b. Continuity equation

For a volume element within the microbial colony, the averaged conservation of mass [39] for the active phase and the aqueous phase is expressed as,

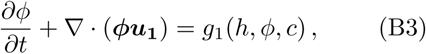

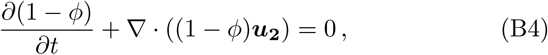

where *φ* is the volume fraction of the active phase (*i.e.* swarmer cells or biomass), ***u*_1_** = (*u*_1_*, w*_1_) is the averaged velocity of the active phase,***u*_2_** = (*u*_2_*, w*_2_) is the averaged velocity of the fluid phase along the *x* (horizontal) and *z* (vertical) directions respectively. Integrating Eqs. (B3) and (B4) through the colony thickness leads to,

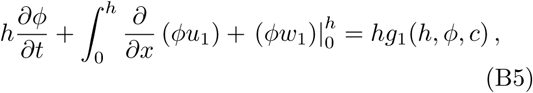

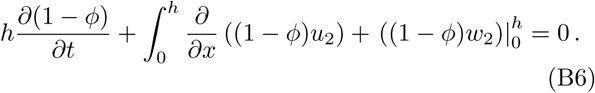

The boundary conditions in Eqs. (B5) and (B6) for the vertical velocities *w*_1_ and *w*_2_ are,

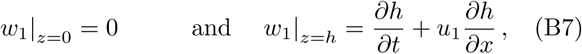

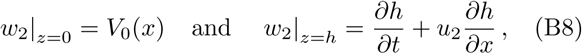

where the no-flux condition is applied to active phase at *z* = 0. For the aqueous phase, at *z* = 0, there is a spatial fluid influx described by *V*_0_(*x*) due to the osmotic pressure difference between the substrate and the microbial colony. Both phases at the upper interface at *z* = *h* obey the kinematic boundary condition. In Eqs. (B2), (B5) and (B6), the 5 unknown fields are *h*, *φ*, *c*, *u*_1_ and *u*_2_. The remaining two equations for closure are obtained from an averaged momentum balance.

### c. Momentum balance

The averaged momentum balances for each phase can be expressed as [38, 39],

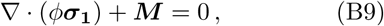

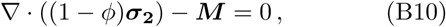

where ***σ*_1_** is the averaged stress tensor acting on a volume element of the swarm cell/EPS matrix phase, ***σ*_2_** is the averaged stress tensor in a volume element of the fluid phase and ***M*** denotes the total momentum transfer between the swarm cell/EPS matrix and the fluid phase. The interfacial momentum transfer term is expressed as [38, 39],

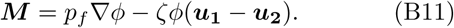

The first term in Eq. (B11) denotes the force *p_f_*∇*φ* due to the averaged interfacial pressure [38, 39] on the cells/EPS matrix by the surrounding fluid. Note that the momentum transfer terms in Eqs. (B9)-(B10) balance each other as the averaged pressures at the two-phase interface are equal, leading to a vanishing net buoyancy. The second term corresponds to the net viscous stokes drag, where *ζ* is a friction coefficient. While Eqs. (B9)-(B11) are generally applicable in describing both swarms and films, particular expressions that describe ***σ*_1_**, ***σ*_2_** and *ζ* are unique to swarming and biofilm expansion and are discussed below for each case.

### 1. Microbial Swarms

Within microbial swarms, the fluid phase is modeled as a Newtonian liquid with viscosity *µ*_2_ whereas the swarmer cells are treated as inviscid with an isotropic stress equal to the surrounding fluid pressure *p_f_*. The averaged constitutive laws for the swarmer cell phase and the fluid phase are expressed as,

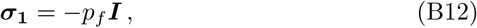

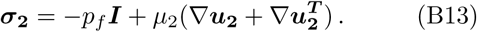

Substituting Eqs. (B12) and (B13) in Eqs. (B9)-(B11), we obtain

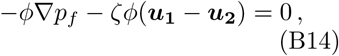

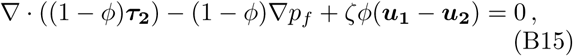

where 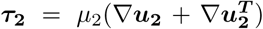 is the deviatoric stress tensor. In bacterial swarms, the friction coefficient is *ζ* = *ζ_c_/V_c_* ≈ 18*µ*_2_*/a*^2^ where *a* is the diameter of the cell, *ζ_c_* ≈ 3*πµ*_2_*a* is the friction coefficient of a single swarmer cell, *V_c_* = *πa*^3^*/*6 is the cell volume.

#### a. Thin-film lubrication limit

We consider the limit of *h* ≪ *R*, where *R* is the colony radius. In the thin-film lubrication limit, combining Eqs. (B14) and (B15) results in the equation governing the mean horizontal fluid velocity to leading order,

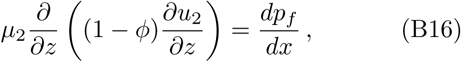

where the fluid pressure is assumed to vary only in the horizontal direction, and is set by the local curvature of the swarm colony and the fluid surface tension as,

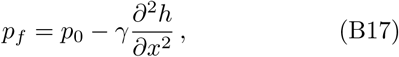

where *p*_0_ is a constant (atmospheric) pressure. Integrating Eq. (B16) twice and using the boundary conditions *u*_2_(0) = 0 and (*du*_2_*/dz*)*_z_*_=*h*_ = 0 lead to the expression for the averaged horizontal fluid phase velocity profile as,

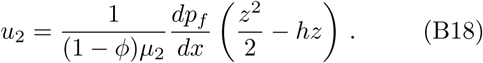

The mean horizontal swarmer cell velocity is determined from the Darcy-type equation in Eq. (B14) as,

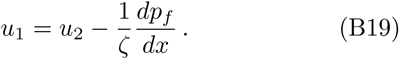

#### b. Growth rate of bacteria in swarms

We note that bacterial swarming is typically associated with nutrient rich environments, where *c* ≫ *K* and *c* ≈ *c*_0_, the initial concentration level. Paradoxically, although the rate of bacterial cell division and volume expansion are exponential, the rate of swarm expansion is constant. Therefore, despite abundant nutrient availability, cell division must be halted at the swarm interior through non-nutrient meditated regulatory/signaling mechanisms, such that only a subpopulation of swarmer cells undergo cell division. In our study, we account for this effect by considering a simple logistic model for the growth term of the form,

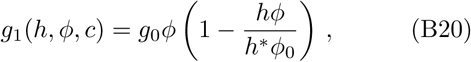

where *h*^∗^ is the swarm colony thickness at the interior, and *g*_0_ is an effective growth rate that accounts for spatial localization in cell division. More specifically, if *L*(*x*) describes the spatial profile of cell growth within a swarm colony, then 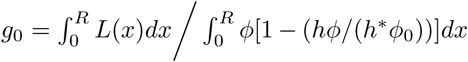, where *R* is the radius of the swarm colony. Measurements of the spatial distribution of cell growth rates within the colony during swarming remain lacking. Consequently, in our model, we determine the value of *g*_0_ by fitting it to the experimental data. We use a nonlinear least-squares solver to match steady-state expansion speeds obtained from solving Eqs. (B21)-(B22) to the experimental data for steady-state *B. subtilis* swarms (see Fig. 1D in main text).

#### c. Swarm equations

Combining Eqs. (B5)-(B8) with Eqs. (B17)-(B19) results in the dimensional thickness averaged equations for swarm colonies,

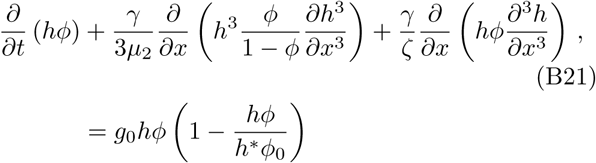

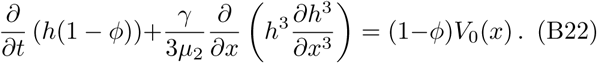

The five boundary conditions are (*∂h/∂x*)*_x_*_=0_ = (*∂h/∂x*)*_x_*_→∞_ = 0, (*∂*^3^*h/∂x*^3^)*_x_*_=0_ = (*∂*^3^*h/∂x*^3^)*_x_*_→∞_ = 0, and *ø*(0) = *ø*_0_, where *φ*_0_ is the volume fraction of the swarmer cells at the interior where there is no flow. As discussed in the main text, we impose the far-field boundary condition by choosing a domain much larger than the colony size.

### 2. Bacterial Biofilms

In our model, the EPS matrix constitutes an active viscous hydrogel network with a polymer volume fraction *φ*(*x, t*), while the fluid phase is considered as a freely moving solvent of volume fraction 1 − *φ*(*x, t*). The averaged constitutive laws for the EPS matrix phase and the fluid phase are expressed as,

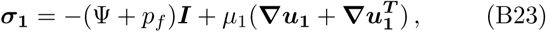

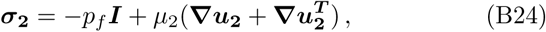

where *µ*_1_ is the viscosity of the EPS matrix, *µ*_2_ is the aqueous phase viscosity, Ψ represents an effective swelling pressure in the biofilm EPS hydrogel and *p_f_* is the fluid pressure. Substituting Eqs. (B23) and (B24) in Eqs. (B9)-(B11), we obtain

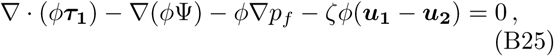

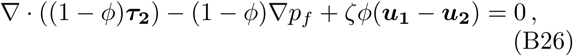

where 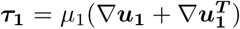 and 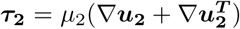 are the deviatoric stresses in the EPS phase and aqueous phase, respectively. In our model, we assume that the contribution of stress from the viscosity of the aqueous phase is negligible compared to the frictional drag due to flow between water and EPS polymer chain network. Consequently, Eq. (B26) reduces to a Darcy-type law of the form,

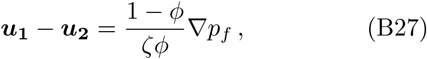

where *κ*(*φ*) = (1 − *ø*)*/ø* is a volume fraction dependent permeability and the fricton coefficient is *ζ* = *µ*_2_*/ξ*^2^, provided that *ξ* ~ 50 nm is the polymer mesh network length scale. Substituting Eq. (B27) in Eq. (B25) results in,

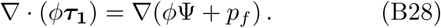

#### a. Thin-film lubrication limit for biofilms

In the thin-film lubrication limit for biofilms when *h* ≪ *R*, Eq. (B28) reduces to

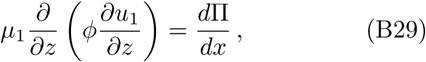

where Π = *φ*Ψ + *p_f_* is treated as an effective EPS phase pressure, as it is *d*Π*/dx* that drives the viscous EPS flow. In Eq. (B29), the effective pressure is assumed to vary only in the horizontal direction, and the fluid pressure is once again set by the curvature and fluid surface tension according to Eq. (B17). Moreover, the swelling pressure is expressed as Ψ = *ψ*(*ø*)*/ø*, where *ψ*(*φ*) = *ψ*_0_ × *φ*^3^ is the osmotic pressure of the biofilm EPS matrix using the Flory-Huggins model for a polymer network in a *θ*-solvent [34, 48]. The osmotic pressure scale is *ψ*_0_ = *kT /b*^3^, where *b* ~ 0.5 nm is the approximate size of the monomer unit in the EPS matrix. Note that although we refer to Π as an effective pressure, in Eq. (B23) the mechanical pressure acting on the EPS phase is instead *ψ*(*ø*)*/ø* + *p_f_*. Integrating Eq. (B29) twice using the boundary conditions *u*_1_(0) = 0 and (*du*_1_*/dz*)*_z_*_=*h*_ = 0 results in,

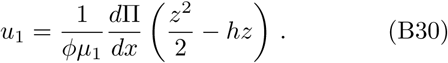

The mean horizontal fluid velocity is determined from the Darcy-type equation in Eq. (B27) as,

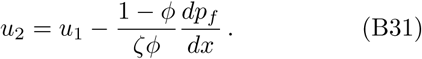

#### b. Growth rate term in biofilms

In biofilms, the growth term *g*_1_(*h, φ, c*) is expressed by the modified logistic term,

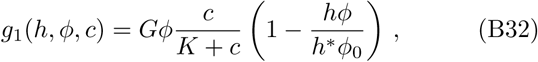

where *G* is the specific EPS production rate, *h*^∗^ is the maximum overall biofilm colony thickness and *φ*_0_ is the volume fraction of the EPS matrix in the fully swollen state. Note that, in contrast to microbial swarming, the growth rate term in expanding biofilms is strongly coupled to the local nutrient concentration field via a Monod-type term in Eq. (B32).

#### c. Biofilm equations

Combining Eqs. (B5)-(B8) with Eqs. (B30)-(B31) results in the dimensional thin-film governing equations for biofilm colonies,

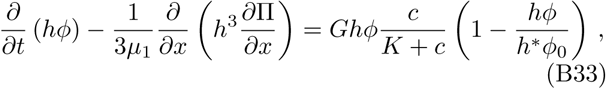

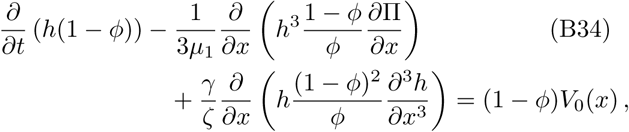

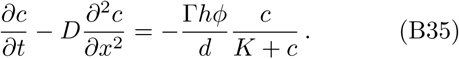

The eight boundary conditions are the symmetry conditions (*∂h/∂x*)*_x_*_=0_ = (*∂h/∂x*)*_x_*_=*R*_ = 0, (*∂h*^3^*/∂x*^3^)*_x_*_=0_ = (*∂*^3^*h/∂x*^3^)*_x_*_=*R*_ = 0, (*∂φ/∂x*)*_x_*_=0_ = (*∂φ/∂x*)*_x_*_=*R*_ = 0, and (*∂c/∂x*)*_x_*_=0_ = (*∂c/∂x*)*_x_*_=*R*_ = 0 where *R* is now the size of the petri dish that is infused with nutrients.

## Appendix C: Numerical computation

Numerical solutions to Eqs. (B21)-(B22) and Eqs. (B33)-(B35) were implemented using the COMSOL 5.0 finite element package. We use a fixed 1D domain of size *x* ∊ [0*, L*_1_] where *L*_1_ = 16 and 150 for the biofilm and swarm simulations respectively. We use quintic Lagrange basis functions with element sizes below 0.04 and use the general form PDE solver. To handle the moving contact line, we introduce a precursor film of thickness *h_p_/h*^∗^ = 0.0125, where *h*^∗^ is the vertical length scale (see Fig. C.1). We follow the regularization described in Ref. [35] to introduce a minimum threshold for growth and a stable fixed point in the precursor film. Specifically, the growth terms in (B2), (B20) and (B32) are multiplied by a factor 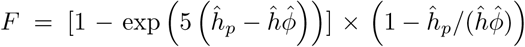, where *F* ≈ 1 everywhere except near the precursor film where *F* = 0.

## ACKNOWLEDGMENTS

We thank H. Berg, L. Ping and A. Pahlavan for helpful discussions. We thank M. Cabeen, R. Losick for providing strains. We thank S. Rubinstein for access to the fluorescent imaging and microbial culture facility. This work was supported by the Harvard MRSEC DMR 1420570 and the MacArthur Foundation (L. M.).

---

**Figure C.1.**
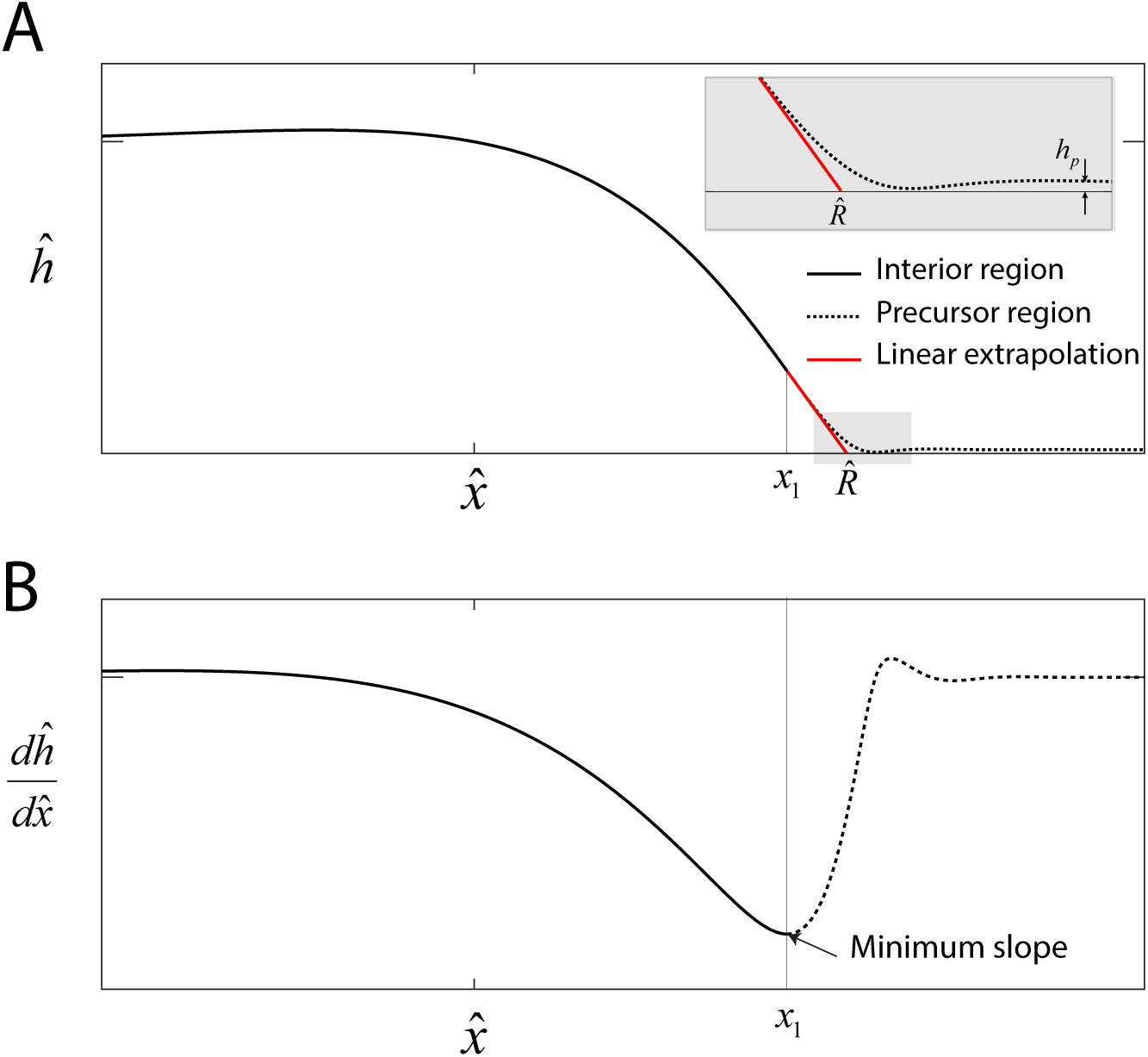
Precursor film. (A) Numerical solution to Eqs. (5)-(6) in the main text that represents the swarm profile 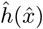. The solid black line indicates the swarm profile in the interior region domain 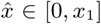. The dashed line beyond *x*1 denotes the precursor-film region. The solid red line represents a linear extrapolation of the swarm profile in the region 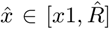, where 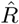 is the swarm radius. Inset: Magnified view of the transition region, where *h_p_* is the precursor film of thickness. (B) The onset of the precursor film is defined at the point of the numerical profile where the slope is a minimum.

**Figure C.2.**
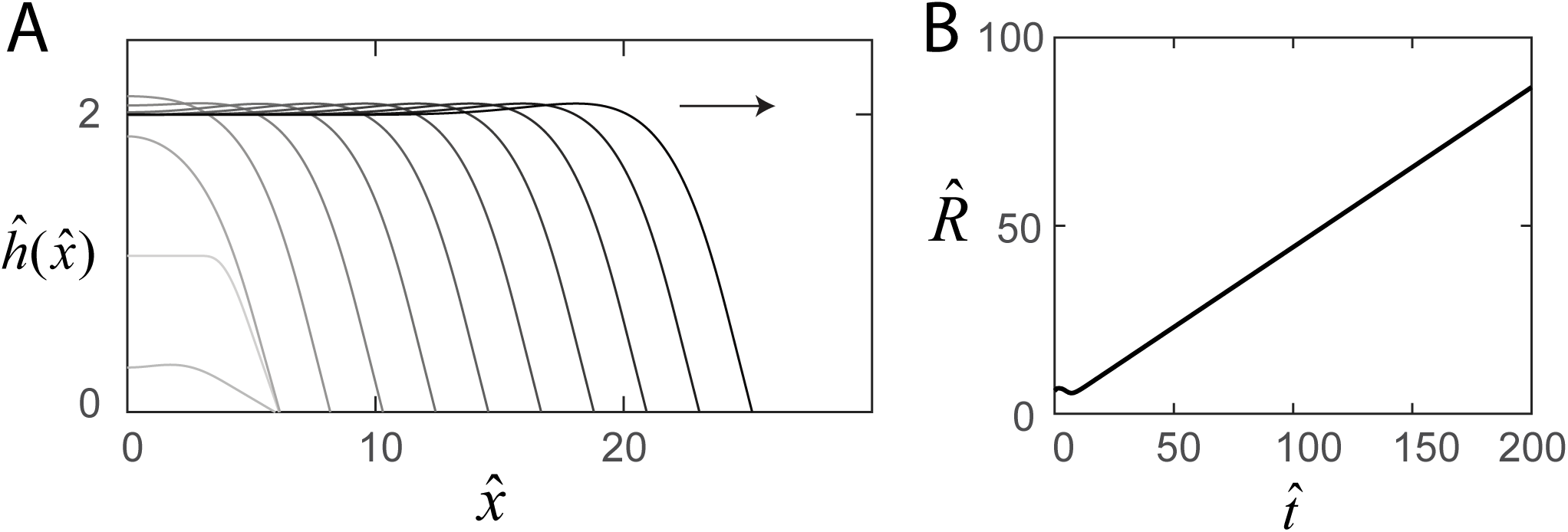
Steady-state swarm solutions. (A) The evolution of the numerical swarm thickness 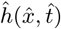 plotted in the laboratory frame at fixed time intervals. (B) Plot of the swarm radius as a function of time indicating steady-state solutions.

**Figure C.3.**
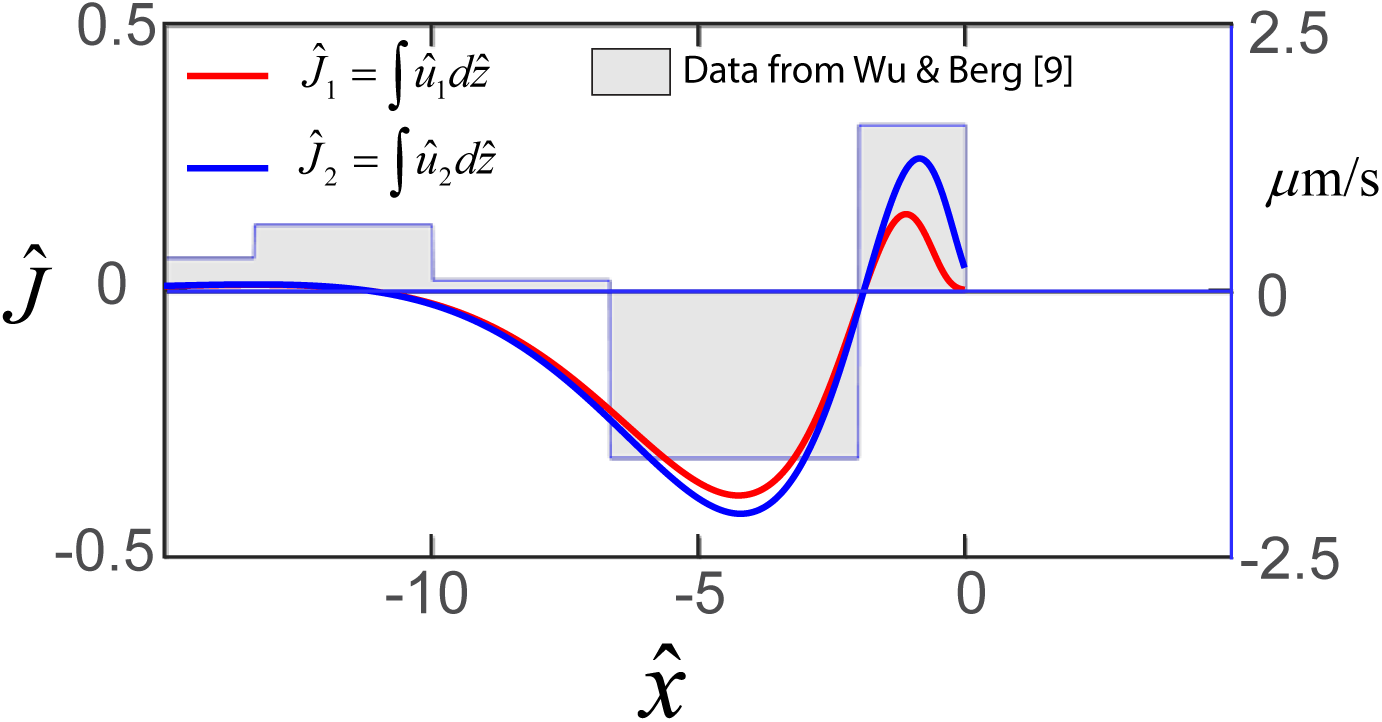
Horizontal flows in swarms. Steady-state profile of the dimensionless net horizontal velocity 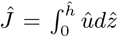 in the swarmer cell phase (red) and fluid phase (blue). Expression for 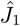 and 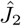are obtained from Eqs. (B18) and (B19) as 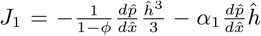 and 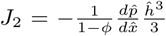 where *α*_1_ = 0.2 as discussed in the main text. The step function represents experimental flow speeds as a function of the distance form the swarm edge as measured by Wu & Berg [22], and where the horizontal distance has been scaled by *L* = 15*µ*m.

**Figure C.4.**
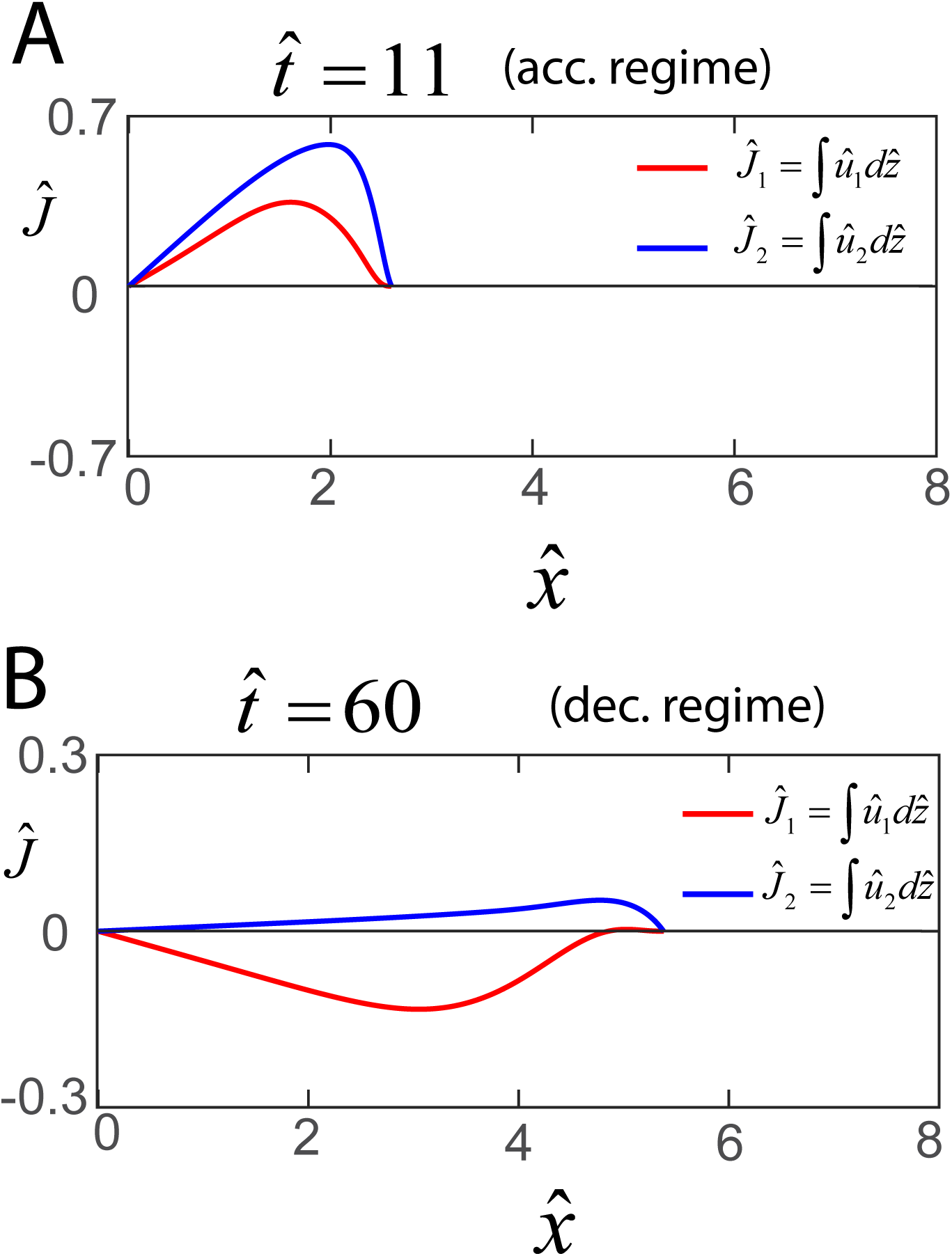
Early and late flows in biofilms. Profile of the dimensionless net horizontal velocity 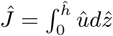 in the swarmer cell phase (red) and fluid phase (blue).</br>Expression for 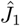 and 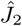 are obtained from Eqs. (B30) and (B31) as 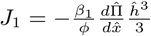 and 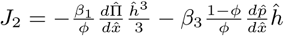, where *β*_1_ = 6.7 and *β*_3_ = 0.02 as discussed in the main text.

